# Actin Branching Regulates Cell Spreading and Force on Talin, but not Activation of YAP

**DOI:** 10.1101/2025.05.09.653153

**Authors:** Claudia Villalobos, Amir Sadeghifar, Jose Maggiorani, Juliet Delapena, Garrett McDaniel, Tristan P. Driscoll

## Abstract

Cells sense the mechanical properties of their environment through physical engagement and spreading, with high stiffness driving nuclear translocation of the mechanosensitive transcription factor YAP. Restriction of cell spread area or environmental stiffness both inhibit YAP activation and nuclear translocation. The Arp2/3 complex plays a critical role in polymerization of branched actin networks that drive cell spreading, protrusion, and migration. While YAP activation has been closely linked to cellular spreading, the specific role of actin branching in force buildup and YAP activation is unclear. To assess the role of actin branching in this process, we measured cell spreading, YAP nuclear translocation, force on the adhesion adaptor protein Talin (FRET tension sensor), and extracellular forces (traction force microscopy, TFM) in 3T3 cells with and without inhibition of actin branching. The results indicate that YAP activation still occurs when actin branching and cell spreading is reduced. Interestingly, while actin de-branching resulted in decreased force on talin, relatively little change in average traction stress was observed, highlighting the distinct difference between molecular level and cellular level force regulation of YAP. While cell spreading is a driver of YAP nuclear translocation, this is likely through indirect effects. Changes in cell spreading induced by actin branching inhibition do not significantly perturb YAP activation. Additionally, this work provides evidence that focal adhesion molecular forces are not a direct regulator of YAP activation.

## Introduction

Integrin-based adhesions form the basis for extracellular matrix mechanosensing and remodeling^1^. However, extracellular matrix (ECM) forces are also directly transmitted through the force generating actomyosin cytoskeleton to the nuclear envelope via the linker of nucleoskeleton and cytoskeleton (LINC)^2^ complex (nesprin, sun, lamin)^3,4^. One mechanosensitive pathway regulated by these forces is the YAP pathway (yes associated protein), which plays a critical role in development, homeostasis, and disease^5–8^.

On 2D surfaces, cell spreading and migration is driven by a branched actin cytoskeletal network that is rapidly polymerizing at the leading edge of the cell to drive rearward flow of actin filaments which can engage and disengage from the integrin receptors embedded in the cell membrane and connected to the ECM. The extent of this engagement and the reinforcement of these connections is dependent on force buildup in force sensitive adapter proteins (talin and vinculin) that directly connect integrins to the moving actin cytoskeleton^9–12^. This cell spreading has been shown previously to be a critical regulator of activation of the mechanosensitive transcription factor YAP^13^, where loss of the focal adhesion protein talin prevents spreading and YAP activation^14^. While stiffness induced cell spreading does require integrin-based focal adhesions, cytoskeletal forces activated with cell spreading are also transmitted to the nucleus, flattening the nucleus and stretching the nuclear envelope^15,16^.

The primary connection between the nuclear envelope and elements of the cytoskeleton is the LINC complex, allowing for transmission of cytoskeletal forces and nuclear flattening during cell spreading^3,15–19^. Previously, we showed that these LINC complex induced nuclear deformations drive activation of the mechanosensitive YAP/TAZ signaling pathway^19^. More recently, others have shown that this is through alterations in the nuclear transport of YAP through nuclear pores^17^. While it is clear this connection critically links nuclear deformation and YAP signaling, it is unclear if branched actin and forces on integrin-based adhesions regulate YAP activation directly.

A major driving factor for the formation and maturation of focal adhesions is the retrograde flow of actin as it is pulled by myosin and pushed by new polymerization via the Arp2/3 complex^20,21^ and formins^22,23^. The branching induced by Arp2/3 regulates the structure of the actin network and creates a balance between protrusion speed versus stability of the leading edge of the cell^21^, which is also force regulated^24,25^. However, the role of this polymerization in facilitating YAP activation with cell spreading is unclear.

Here, we implemented a variety of perturbations to actin branching that reduce cell spreading to assess their influence on YAP activation, force on the focal adhesion adapter talin, and traction stresses on the extracellular matrix. We observe that YAP activation occurs independent of cell spreading and force on the focal adhesion protein talin. Together, these results indicate that while cell spreading can indirectly regulate YAP through nuclear flattening and pore deformation, cell spread area and talin forces are not direct regulators of YAP activation.

## Methods

### Cell Culture and Transfections

NIH 3T3 cells were cultured on tissue culture dishes at 37°C, 5% CO2 in Dulbecco’s Modified Eagle Medium (DMEM, Invitrogen) with 10% FBS and 1% Penicillin-Streptomycin (Invitrogen) and passaged 1:10 every 3 days. Talin tension sensor (TS) and c-terminal control sensor (CTS) were transiently transfected into NIH 3T3 cells for FRET experiments. All plasmids expression constructs were prepped from chemically competent bacterial cells (GC10a, Genessee Scientific) using the ZymoPURE II Plasmid Midiprep Kit (Zymo Research). DNA transfections were performed using jetPRIME transfection reagent following the manufacturer protocol. Briefly, 2 μg of DNA was diluted in 200 μl of jetPRIME buffer and vortexed for 10 seconds. 4 μl jetPRIME reagent was added, vortexed for 1 second, and incubated at room temperature for 10 minutes. The transfection mix was then added to cells in 2ml of media in a 6- well plate. Cells are incubated at 37°C in transfection mix overnight and then media is replaced with fresh media. Cells were used for experiments 36-72 hours post transfection. For FRET and staining experiments, glass coverslips were coated with fibronectin at a concentration of 10μg/ml in phosphate buffered saline (PBS, pH 7.4) at 4°C overnight. They were rinsed 3 times with 1x PBS to remove excess fibronectin prior to cell seeding.

### Immunostaining

Immunostaining was performed as previously described^26^. The cells were fixed with 4% paraformaldehyde (PFA) for 15 minutes at room temperature. Following fixation, cells were washed three times with 1x PBS and permeabilized for 5 minutes using 0.05% Triton X-100 in PBS supplemented with 320 mM sucrose and 6mM magnesium chloride (cytoskeletal stabilizing buffer). Samples were blocked with 1% w/v BSA in PBS for 30 minutes at room temperature and then stained with primary antibody for YAP (Santa Cruz, c-101199, 1:200 in 1% BSA-PBS). Cells were washed 3 times with PBS and then stained with Alexa-488 phalloidin (Thermo Fisher) and secondary antibody Alexa-568 goat anti-mouse (1:1000, Thermo Fisher) and Alexa-647 goat anti-rabbit (1:1000, Thermo Fisher) antibodies. Cells were washed 3 times with 1x PBS and mounted with Fluoromount G with DAPI (Southern Biotech) to label nuclei. Images were acquired on a Zeiss AxioObserver 7 with Colibri7 light source and Hammamatsu Flash 4.0 sCMOS camera using a 20x 0.8NA objective. Quantification of cell area and nuclear to cytoplasmic ratio for YAP was performed in MATLAB.

A custom MATLAB program was used to detect multiple parameters from immunostained images^27^. Canny edge detection was used to detect the nuclear areas from the DAPI images and generate masks for calculation of the average intensity of the signal in the nucleus. Nuclear mask segmentation was performed using the MATLAB bwlabeln function to group nuclear pixels into individual masks for each nucleus. A second mask was generated by dilating the nuclear area mask to generate a mask of a ring of cytoplasm just outside the nucleus (5-pixel width). This second mask was used to calculate the average cytoplasmic signal and the ratio of nuclear to cytoplasmic signal was calculated for each cell. While this does not average signal over the entire cytoplasm, it provides average signal from a similar thickness area of the cell. ImageJ software was used to quantify the cell area using the F-Actin signal using image thresholding and the analyze particle function.

### FRET Imaging and Analysis

Cells were imaged on Zeiss AxioObserver 7 with 63x 1.4NA objective, Colibri7 epifluorescence light source and Hammamatsu ORCA Flash 4.0v3 camera. 16-bit images were acquired for 3 channels (each with 0.5s exposure). Images were acquired with the following LED/filter combinations: donor (GFP) channel with a 488-nm (excitation LED) and 510/50 filter (emission), acceptor (tagRFP) with 560-nm LED (excitation) and 590/70 filter (emission), and FRET channel with a 488-nm LED (excitation) and 590/70 filter (emission). FRET values were calculated using custom scripts in MATLAB as previously described^37^. All three FRET images (eGFP, FRET, tagRFP) were corrected for illumination gradient, pixel shift, and background subtraction followed by three-point smoothening. Bleed-through and cross-excitation coefficients were calculated by imaging cells transfected with only eGFP-talin or TagRFP-talin and performing a linear regression on the signal (Supplemental Figure 1). The slope of the pixelwise FRET channel intensity versus the donor or acceptor channel intensity gives donor leakage (dL) and acceptor leakage (aL) fractions, respectively. Heat maps of FRET and pixelwise FRET index were calculated using the following equation:

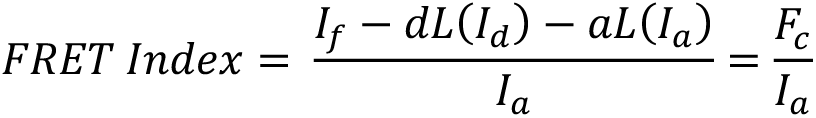

where *I_f_*, *I_d_*, and *I_a_* are the shade, shift, and background-corrected pixel intensities for each of the respective channels (fret, donor, and acceptor) and *F*_$_ is the corrected FRET intensity.

### PDMS and Polyacrylamide Substrate Preparation

PDMS substrates were made using Sylgard 184 PDMS kit (electron microscopy sciences, Catalog #24236-10). PDMS base (B) and curing agent (C) were mixed for 10 minutes to make 2kPa, or 10kPa gels (B/C ratio of 70/1 or 55/1)^28^. The gels were degassed under vacuum to remove bubbles and the mixture was spin coated at approximately 4000 rpm for 30s to achieve a ∼50µm thick layer. The dishes were cured by incubation at 70°C for 3 hours.

Very soft polyacrylamide substrates (0.5kPa) used to completely inhibit cell spreading were fabricated as described previously^29^. Briefly, 20-mm coverslip-bottomed dishes (#0 coverslip; Mattek) were silanized with a 2% solution of 3-aminopropyltrimethoxysilane (APTES) in isopropanol for 10 min at room temperature. After washing with double-distilled H2O (ddH2O) and drying, coverslips were incubated with 1% glutaraldehyde solution in ddH2O for 30 min and then washed three times with ddH20 and dried. Polyacrylamide gels were cast onto the silanized surface by preparing a 3% acrylamide and 0.06% bis-acrylamide solution (Bio-Rad) and polymerizing with ammonium persulfate 0.1% w/v (Sigma) and 0.15% v/w TEMED (Sigma). Gels were cast between the silanized glass surface and a 12-mm uncoated glass coverslip with a volume of 12 μL. After casting, gels were treated with fresh sulfo-SANPAH (Sigma) in ddH2O (2 mg/mL) and exposed to ultraviolet (UV) light for 3 min (8 W, 254-nm wavelength at a distance of 2 in). After UV, gels were washed with ddH2O and then incubated with fibronectin overnight (10μg/mL in PBS at pH 7.4).

### Traction Force Microscopy (TFM)

For TFM experiments, 10kPa PDMS substrates were coated with fluorescent beads as previously described^26,30^. The PDMS surface was exposed to UV light for 30 minutes and then treated with (3-Aminopropyl)triethoxysilane (APTES, Sigma) 2% w/v in isopropanol (Sigma) to functionalize the surface of the gels. Gels were washed 3 times with water and then treated with N-(3-Dimethylaminopropyl)-N’-ethylcarbodiimide hydrochloride (EDC, 100ug/ml, Sigma) in borate buffer solution with fluorescent beads (20mL EDC buffer solution with 7µL of freshly sonicated FluoSpheres carboxylate modified 0.1µm, blue 350nm). Substrates were then washed three times with 1x PBS, UV sterilized and coated with fibronectin (overnight at 4°C, 10 μg/ml). Cells were seeded on TFM substrates and imaged in a live cell imaging chamber under phase contrast (for cell area) and epifluorescence (for bead images) before and after adding sodium dodecyl sulfate (SDS, 0.1% w/v) to lyse the cells and collect both the deformed and undeformed gel bead images. Cell lysis was confirmed by phase contrast after SDS treatment. The TFM images were acquired with an oil immersion objective (63x, 1.4NA) allowing for high resolution imaging of the 100nm fluorescent beads. These images were processed using previously developed MATLAB code^31^ using Fourier Transform Traction Cytometry (FTTC). The gels were assumed to be incompressible with Poisson’s Ratio of 0.5 and the regularization parameter was chosen to be 0.0001 to average out discontinuity errors in the displacement field. Traction force maps were generated to determine the average traction stress for each cell using cell area masks generated in ImageJ based on phase contrast images of the cells acquired before cell lysis.

### mRNA Isolation, cDNA Synthesis, and qPCR

For mRNA analysis with 24 hours of inhibition, mRNA isolation was carried out using the Spin protocol of the Aurum^TM^ Total RNA Mini Kit (Biorad). cDNA synthesis was performed using Biorad iScript^TM^ Reverse Transcription Supermix for RT-qPCR protocol in a Biorad T100^TM^ Thermal Cycler. qPCR was then performed in a 96-well plate in duplicate using the iTaq Universal SYBR^TM^ Green Supermix in the Applied Biosystems^TM^ 7500 rt-PCR instrument and fold change in expression was calculated using the delta delta CT method with normalization to β-Actin. Primers targeted mouse CTGF (fw: 5’-ctgcagactggagaagcaga-3’, rv: 5’-gatgcactttttgcccttctt-3’) and β-Actin (fw: 5’-cgagcgtggctacagcttc-3’, rv: 5’-gccatctcctgctcgaagtc-3’),

### Statistical Analysis

Each experiment was replicated 2-3 times as indicated in the figure legends. Statistical tests for each experiment are indicated in the figure legends and significance was set at p<0.05. Violin plots show distribution, mean, and quartiles. All data were tested for normality using a Shapiro-Wilk test (with alpha=0.05). Normally distributed data were tested using One-Way ANOVA with Tukey’s post hoc. Data that were not normally distributed were tested using a non-parametric Kruskil-Wallis test with Dunn’s post hoc. Bar plots indicate mean +/-standard error of the mean. All plotting and statistical analysis was performed in GraphPad prism 9.

## Results

### 4.1 Inhibition of actin branching reduces cell spreading but does not prevent YAP activation

To assess the influence of actin branching on cellular spreading and activation of the mechanosensitive transcription factor YAP, we inhibited actin branching using a small molecule inhibitor of the Arp2/3 complex (CK666, 50μM) or by inhibition of Rac (NSC23766, 50μM). These inhibitor doses were chosen so that they would limit cell spreading without completely blocking it. 3T3 cells were seeded in the presence of DMSO (control) or inhibitors on fibronectin coated glass and allowed to spread for 2 hours. During this early spreading, cells are heavily dependent on Arp2/3 and Rac to generate lamellipodia. Cells were then fixed with 4% PFA and stained for F-actin for quantification of cell area, DAPI for quantification of nuclear area, and YAP to quantify nuclear localization of YAP. There was a significant decrease in both cell area and nuclear area for the CK666 groups (Figure 1 A-B), consistent with the important role of Arp2/3 in early cell spreading. Interestingly, there was no significant change in nuclear localization of YAP (Figure 1 C-D). As a control, we also performed experiments at this early time point where no spreading was possible (through seeding on very soft 0.5kPa polyacrylamide gels). Consistent with previous work, these methods totally prevented nuclear localization of YAP (Supplemental Figure 2 A-C). These results indicate that normal YAP activation still occurs with reduced cell spreading, but that completely blocking cell spreading prevents activation.

**Figure 1:**
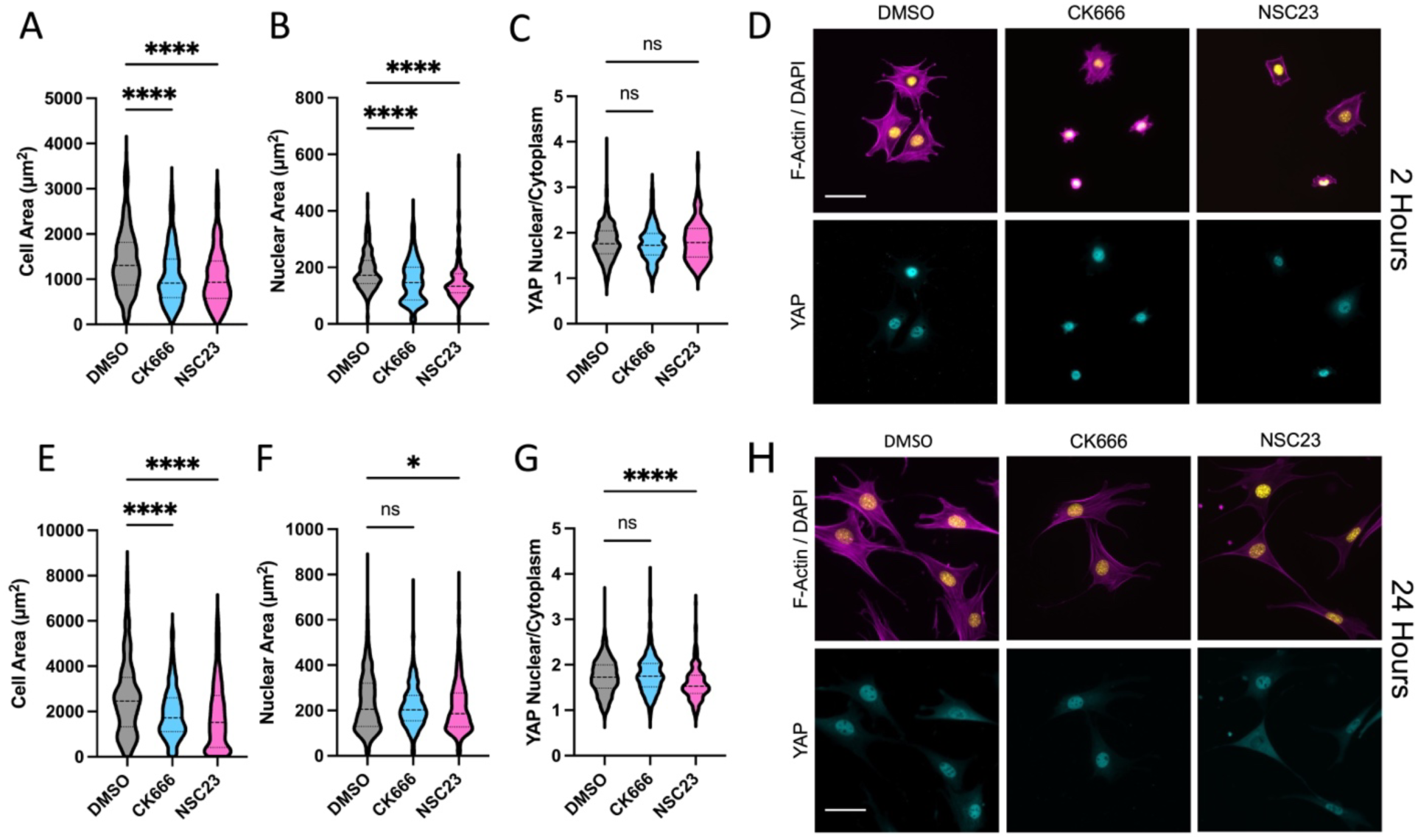
Quantification of cell spread area (A), nuclear area (B), and YAP nuclear localization (C) with reduced actin branching by inhibition of Arp2/3 (CK666 50μM) or Rac (NSC23 50μM) during 3T3 cell spreading for 2 hours on fibronectin coated glass. Representative images of DAPI/F-Actin and YAP (D). Similar quantification for cells allowed to spread for 24 hours with inhibition (E-H). Violin plots indicate distribution, mean, and quartiles, n=215-432 cells/grp from 3 independent experiments. Non-Parametric Kruskal-Wallis test and Dunn’s multiple comparison. (* p<0.05, **** p<0.0001. Scale bar = 50μm.)

Rac GTPase signaling plays an important role in cell spreading, actin branching, and activation of lamellipodial protrusion. As an alternative method of inhibition of spreading and actin branching we inhibited Rac using the small molecule inhibitor NSC23766. As expected, Rac inhibition also resulted in reduced cell spreading and nuclear area (Figure 1 A-B). However, quantification of YAP nuclear localization showed no change in nuclear localization of YAP (Figure 1 C-D). This further indicates that perturbations to actin branching induced cell spreading do not directly regulate YAP activation, and that for partially spread cells, YAP can still be strongly activated.

To assess if these changes persist at longer time points, cells were seeded with or without inhibitors for 24 hours, and the same analysis was performed (Figure 1E-H). Both Arp2/3 inhibition and Rac inhibition resulted in significantly lower cell area. YAP was still nuclear at this longer time point for both groups. Analysis of expression of connective tissue growth factor (CTGF) which is a downstream target gene of YAP, showed no significant change for Arp2/3 or Rac inhibition, but did show a significant reduction for myosin inhibition which is known to regulate YAP (Supplemental Figure 3). However, there was a small but significant reduction in nuclear YAP for the Rac inhibited group, possibly due to crosstalk between Rac signaling and YAP via the canonical Hippo signaling pathway through Lats1/2^32^.

As an alternative method to perturb Arp2/3, we overexpressed a GFP tagged version of the actin debranching protein GMFβ^33^ and performed a similar cell spreading and YAP activation experiment. Consistent with results using CK666, debranching of actin using a separate non-pharmacologic perturbation reduced cell spreading (Figure 2A,B) but did not prevent nuclear spreading and translocation of YAP at early time-points. Instead, this resulted in increased YAP activation (Figure 2C,D), possibly due to changes in actin structure that alter the transmission of forces to the nucleus. However, at later time-points, normal cell spreading, and YAP activation were observed (Supplemental Figure 4). Combined, these results provide support for a YAP mechanical activation mechanism that is not purely dependent on cell spreading.

**Figure 2:**
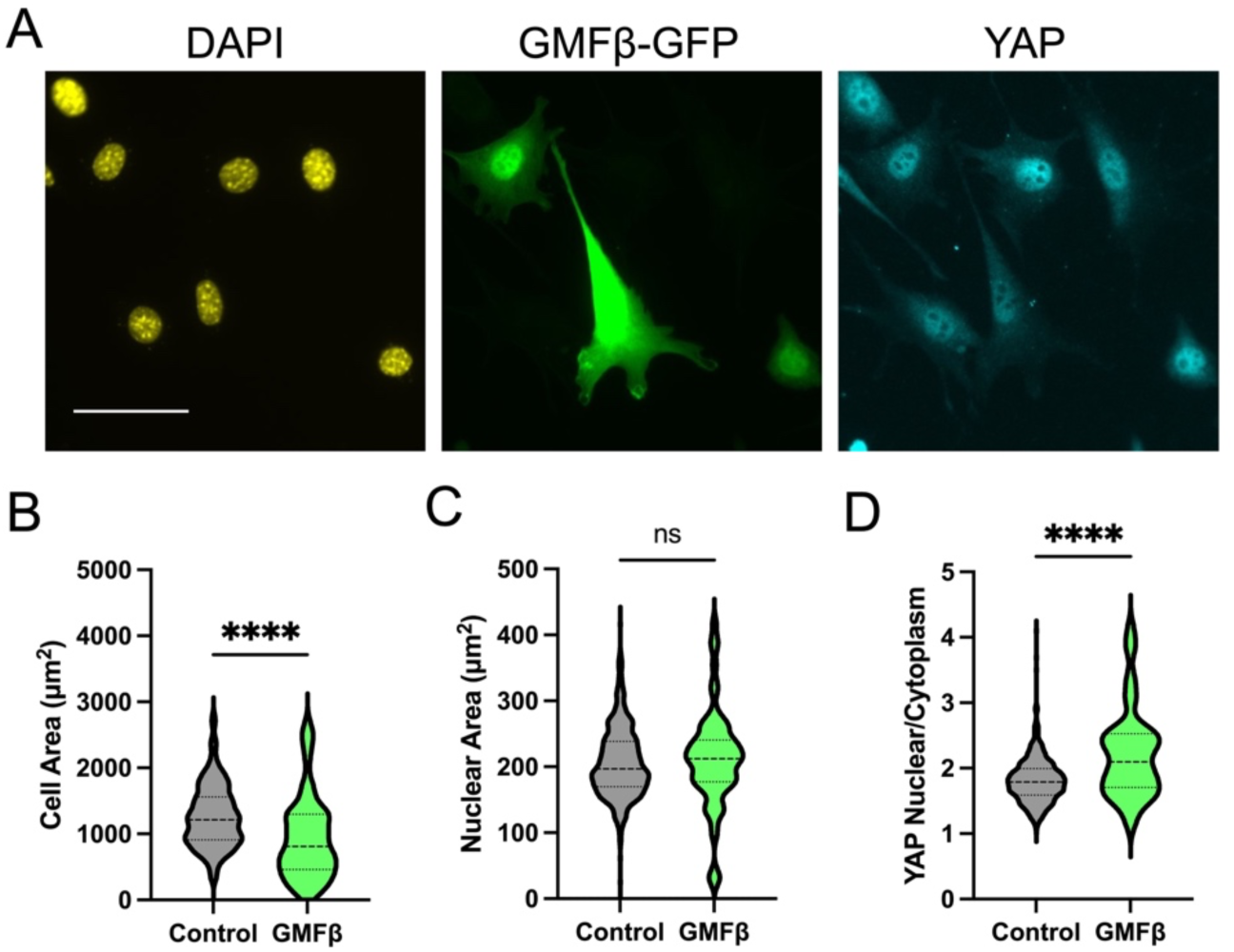
Reducing actin branching by overexpression of the actin debranching protein GMFβ labeled with GFP. Representative images of GMFβ expressing cells spread on fibronectin coated glass for 2 hours (A). Quantification of cell area (B), nuclear area (C), and nuclear to cytoplasmic ratio of YAP (D). Violin plots indicate distribution, mean, and quartiles, n=77-368 cells/group from 2 independent experiments (scale bar = 50μm). Non-Parametric Kruskal-Wallis test and Dunn’s multiple comparison.

### 4.2 FRET Tension Sensor Measurements of Force on Talin

Since the focal adhesion protein talin is the central adaptor and required for cell spreading and YAP activation on high stiffness^14^, we next assessed intracellular forces on talin using a previously described FRET-based talin tension sensor^34,35^. Transfected cells were treated with DMSO or CK666 and compared to a high FRET (no force) control sensor where the sensor module is attached to the c-terminus of talin (Figure 3A). Arp2/3 inhibited cells showed a reduced cell spread area as before. Analysis of the FRET index in talin focal adhesion within these cells showed a significant increase in average FRET compared to DMSO treated control, indicating a decrease in force on talin (Figure 3B). Control sensor showed high FRET (low force) as expected (Figure 3A, CTS). Combined, this indicates that the debranching of actin and reduced spreading that occurs with Arp2/3 inhibition reduces the average force per talin molecule within focal adhesions. Importantly, these FRET measurements are only analyzed in focal adhesions and normalized to adhesion area. Thus, these changes in FRET indicate alterations in the average force per molecule in the focal adhesions.

**Figure 3:**
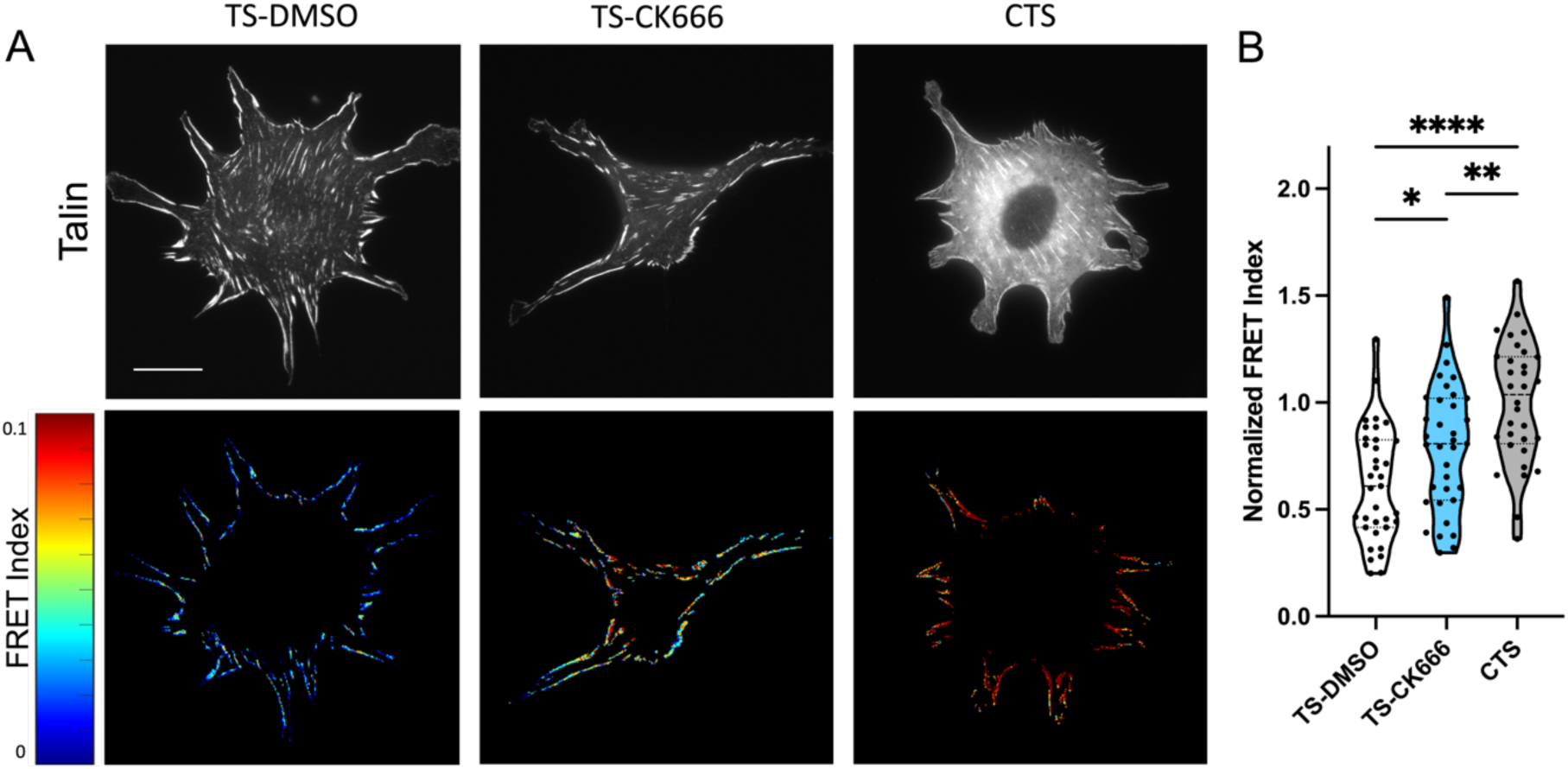
(A) GFP images and FRET index heatmaps of talin tension sensor (TS) or control sensor (CTS) expressing cells treated with DMSO or CK666 inhibitor on fibronectin coated glass. (B) Average FRET index per cell. (Violin plot indicates distribution, mean, and quartiles, n=32-35 cells/group from 3 independent experiments, * p<0.05, *** p<0.001, **** p<0.0001, One-way ANOVA with Tukey’s multiple comparison, Scale bar = 50μm)

### 4.3 Deformable Substrates and Traction Force Microscopy

Cellular spreading results in physical engagement with the substrate and cellular induced traction stresses, regulating YAP activation in a substrate stiffness dependent manner^13^. We next looked to assess the role of Arp2/3 in extracellular forces using traction force microscopy (TFM). Since these measurements require quantification of cell induced deformations in a deformable substrate, we first repeated the previous experiments on PDMS substrates that can be used for TFM measurements. To assess each of these metrics all on a single stiffness, we chose a moderate stiffness PDMS gel (10kPa) that was stiff enough to activate cell spreading, focal adhesion formation, and YAP nuclear translocation, but soft enough to calculate stress fields from bead displacements. As an additional control, we included the myosin inhibitor blebbistatin, which blocks cellular contractile forces but does not prevent stiffness dependent cellular spreading^36^.

Cells were seeded on fibronectin coated 10kPA substrates and allowed to spread in the presence of DMSO (control), CK666 (50μM), or the myosin inhibitor Blebbistatin (10μM) (Figure 4A). Consistent with glass experiments, inhibition of Arp2/3 significantly reduced cell spreading (Figure 4B) but did not reduce nuclear area (Figure 4C) or prevent activation of YAP (Figure 4D). Inhibition of myosin with Blebbistatin did not reduce cell spreading, instead resulting in a slight increase in cell spread area (Figure 4B). Despite this increased cell spreading, there was a significant reduction in YAP nuclear localization with inhibition of myosin contractile forces (Figure 4D).

**Figure 4:**
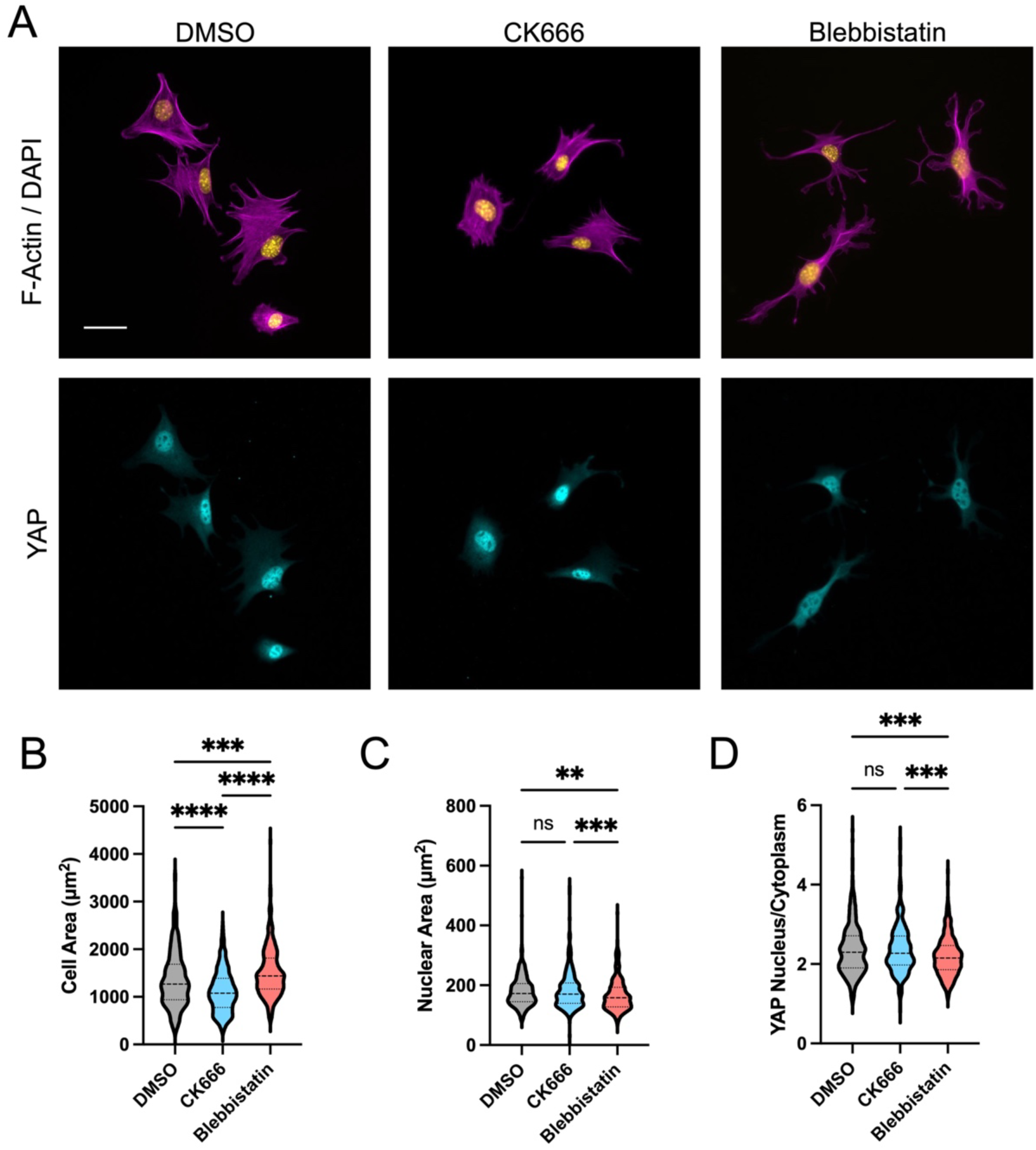
Cell spreading on deformable 10kPa fibronectin coated PDMS substrates with inhibition of Arp2/3 (CK666, 50uM) or myosin (Blebbistatin, 50uM). Quantification of cell area (B), nuclear area (C), and YAP nuclear to cytoplasmic ratio (D) (Violin plots indicate distribution, mean, and quartiles, n=360-422 cells/group from 3 independent experiments, Non-Parametric Kruskal-Wallis test and Dunn’s multiple comparison).

Next, we assessed traction forces and adhesion forces on these deformable PDMS substrates. Average traction stress on this moderate stiffness were relatively unperturbed with CK666 (Figure 5A,B). The blebbistatin group showed close to zero average stress as expected, since myosin is responsible for the vast majority of extracellular traction (Figure 5B). Similar results were also observed for both CK666 and NSC on softer gels (Supplemental Figure 5). Analysis of talin forces on 10kPa gels showed a significant reduction in force on talin with Arp2/3 inhibition (Figure 5C,D). This is consistent with the previous experiments on fibronectin coated glass, indicating that loss of branching does reduce the force on talin and cell spread area without significantly impacting the activation of YAP. Interestingly, the average traction stress is also relatively unperturbed, indicating that the force exerted per cell area and how it is propagated through the cytoskeletal network is a more important regulator of YAP than the force on individual talin molecules.

**Figure 5:**
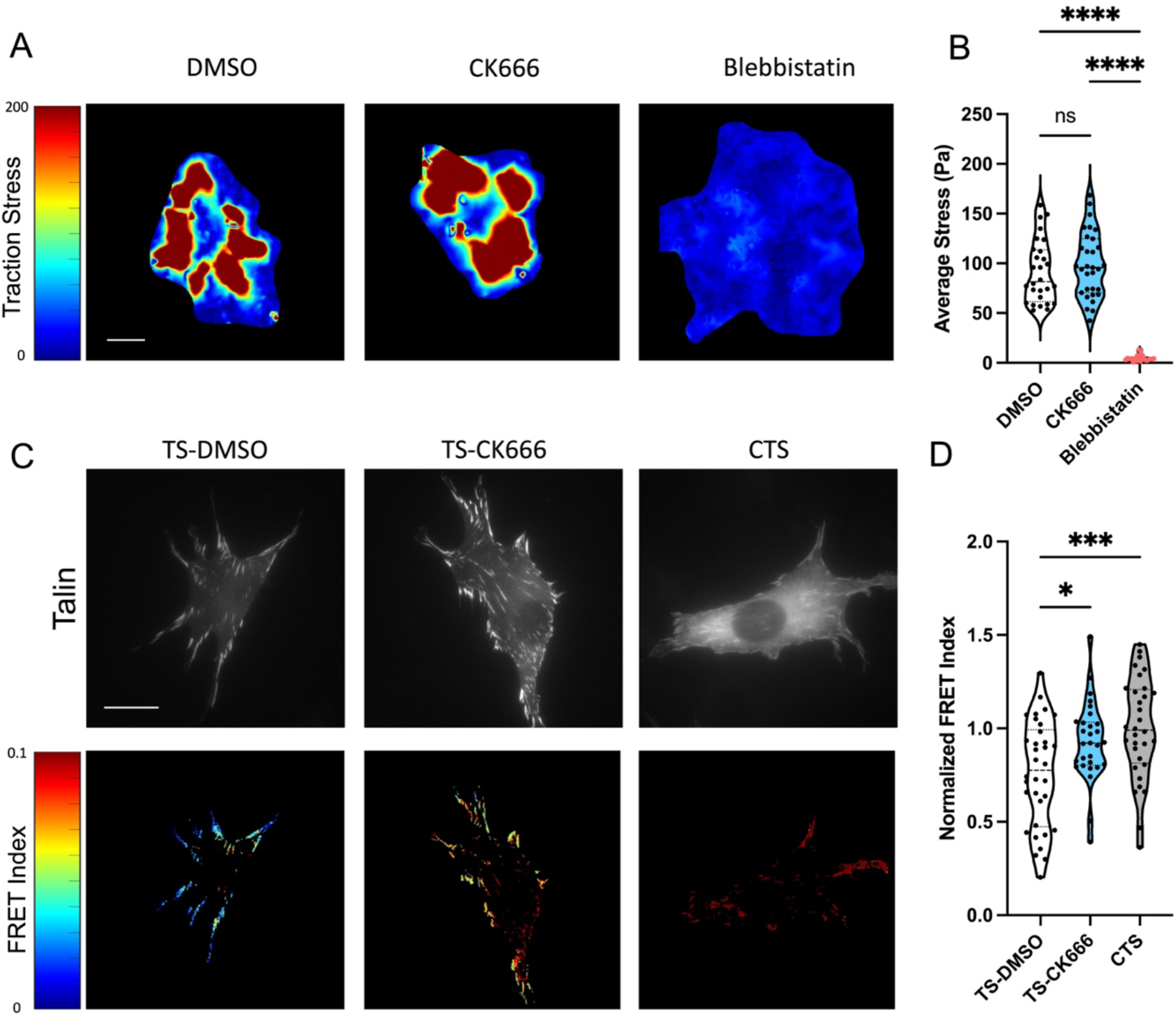
Arp2/3 Inhibition on moderate stiffness (10kPa) alters force on the focal adhesion adapter protein talin. Traction force microscopy for cells seeded on 10kPa PDMS gels and treated with CK666 or Blebbistatin for 2 hours during spreading. (A) Average traction stress per cell and (B) representative traction stress fields. Violin plot indicates distribution, mean, and quartiles. One way ANOVA with Tukey’s post hoc. n=29-32 cells per groups from 2 independent experiments. **** p<0.0001. Representative images and FRET heatmaps for 3T3 cells expressing Talin seeded on 10kPA PDMS substrates and normalized FRET index (normalized to C-terminal Control sensor (CTS)) (D) (Violin plots indicate distribution, mean, and quartiles, n=29-34 cells, from 2 independent representative experiments). One way ANOVA with Tukey’s post hoc. * p<0.05, *** p<0.001, **** p<0. 0001.

## Discussion

The regulation of YAP, and its ortholog TAZ, was originally described as being primarily though the canonical Hippo pathway that is essential for organ size control in both mammalian and non-mammalian systems^37,38^. In canonical Hippo signaling, cell-cell contact results in activation of LATS1 and LATS2 kinase, which phosphorylates YAP to sequester it in the cytoplasm. Loss of cell-cell contact would then activate YAP through inactivation of the Hippo pathway, allowing for YAP to translocate to the nucleus to associate with TEAD and transcribe target genes. Later work by Dupont and Piccolo showed that a second major driver of YAP regulation is independent the canonical Hippo pathway and LATS1/2. This work showed very clearly that inhibition of cell spreading through restriction of substrate area using micropatterned surfaces could completely inhibit the activation of YAP. Further, they showed that this activation is independent of LATS1/2, but instead regulated by an F-Actin / cytoskeletal force related mechanism. Following this work, much of the literature has emphasized the importance of cell spreading and cell spread area on the regulation of YAP. Inherently, cell spread area is directly regulated by a cell’s ability to make physical connections with its extracellular environment using integrin-based adhesions. More recent work has shown that deformation of the nuclear envelope plays a key role in the regulation of YAP, and that this regulation can occur independent of focal adhesions and cell spreading^17^. However, the direct connection that the cytoskeleton provides between focal adhesions and the nuclear envelope through the LINC complex makes adhesions critical in the network level force transfer that is required for these nuclear deformations.

Talin is essential for integrin-based adhesions, since it is the central adapter molecule that directly connects integrin to the actin cytoskeleton^39,40^. As such, loss of talin completely prevents focal adhesion formation, inhibiting sustained cellular spreading^41^. As expected, this also prevents YAP activation due to the severe adhesion and cell spreading defect that results from the loss of this essential focal adhesion protein^14^. Talin is also essential for mechanosensing of extracellular matrix stiffness, often attributed to its ability to undergo force dependent unfolding events to recruit vinculin and stabilize adhesions^10,42^. We have previously shown that talin displays increasing force per molecule with increasing matrix stiffness^34,35^.

Here, we identify that the magnitude of this force per molecule is not a direct regulator of mechanosensing through YAP. Despite a significant change in the force per talin molecule with actin de-branching, there is relatively little impact on the activation of YAP. Previous work has also shown that YAP activation itself can feedback to regulate adhesion stability and influence adhesion molecule gene expression^43^. Loss of YAP or YAP activity can also inhibit spreading^44^. Importantly, we have studied these impacts on talin force at very early time points (2hrs), to avoid the impacts of feedback from YAP to influence adhesion stability and adhesion molecule expression levels.

Actin polymerization has been very clearly shown to be essential for YAP/TAZ activation.

Inhibition of actin polymerization with Latrunculin A results in complete inhibition of YAP^45^. However, this also completely prevents cell spreading, focal adhesion formation, and actin induced nuclear deformations, making the interpretation convoluted. While the inhibition of myosin does also reduce YAP activation, this inhibition is incomplete. Despite a significant loss of focal adhesion formation with myosin inhibition, cell spread area and stiffness dependent cell spreading behaviors are maintained^36^, and YAP is still localized to the nucleus on high stiffness but does show a slight reduction in fibroblasts^45^. Here, we target just one aspect of actin polymerization (branching based polymerization) and observe that despite a significant impact on cell spreading, the nuclear localization of YAP is unperturbed. This is also consistent with previous work that has shown bundled actin to be more important than branched actin for the induction of the YAP target gene CTGF^46^. Therefore, while branched actin polymerization and focal adhesion formation is required for normal cell spreading, it is not required for activation of YAP.

While it may seem counter intuitive that the force per talin molecule could be reduced while the force on the extracellular surroundings could be maintained, this discrepancy could be explained by the degree of force coordination. With changes in the distribution of force orientations at the molecular level, the force in individual adhesion molecules (talin tension sensor) can be decoupled from force at the scale of the cell (traction forces) (Figure 6). In normal cells that have a high degree of actin branching, forces acting on the adhesion molecules are high, but the branching allows for force in multiple directions that are potentially uncoordinated (Figure 6A). With de-branching, these forces become coordinated, such that lower force per molecule is required to generate a high coordinated traction stress (Figure 6B). At the cellular level, cells with branched actin have high cell area, high traction, and activation of YAP on substrates of medium and high stiffness (Figure 6C). With de-branching, the cell area is reduced, but normal transmission of traction stress and normal activation of YAP (Figure 6D) are maintained. This is potentially due to the more coordinated forces in adhesion molecules.

**Figure 6:**
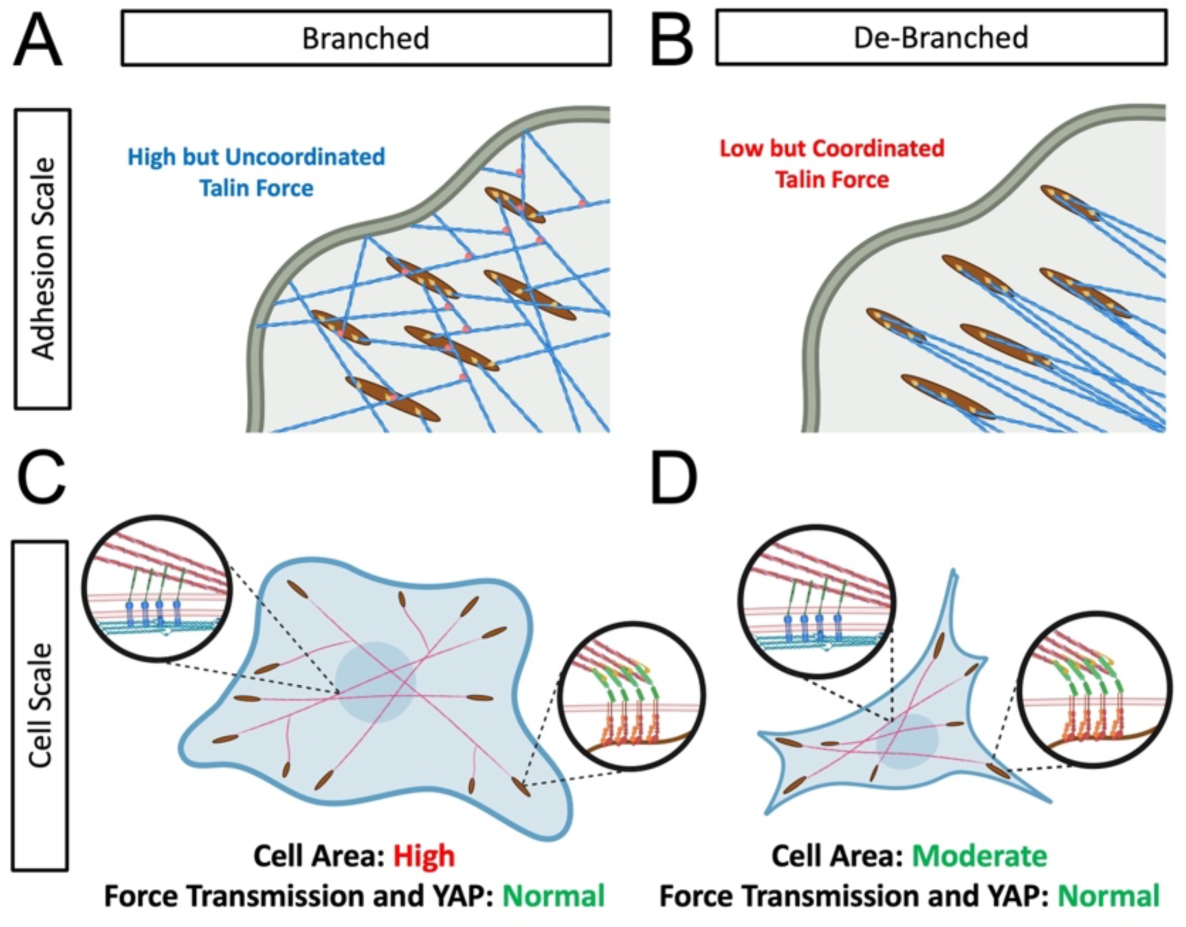
Schematic illustration of the impact of de-branching on the adhesion scale force distribution and the cell scale force distribution. At the scale of individual adhesions (A) branched actin networks (A) result in high forc on talin that is likely less coordinated. De-branching reduces the force per talin molecule, but these more coordinated talin forces are still able to generate traction on the ECM (B). This results in cellular scale force that are normal (C) for cells regardless of branching (D), and still maintains force in the actin cytoskeleton necessary to deform the nuclear envelope and activate YAP/TAZ mechanosensing to normal levels.

However, this could also indicate alterations in the structure of the focal adhesions or alterations in the recruitment of other focal adhesion molecules such as vinculin, which is known to directly bind Arp2/3, recruiting it to sites of integrin clustering^47^. In both cases, contractile actin forces that deform the nucleus are maintained, allowing for normal activation of YAP through the previously described mechanism of LINC complex induced nuclear deformation^17,19^.

More targeted approaches to assess the impact of focal adhesion proteins and integrin signaling in the regulation of YAP, have also indicated a role for vinculin, talin, and focal adhesion kinase (FAK)^14,48,49^. Loss of vinculin, loss of talin, or mutation of vinculin binding to talin all result in reduced nuclear area as well as the nuclear localization of YAP^48^. Furthermore, inhibition of FAK, loss of FAK, loss of FAK catalytic activity, FAK activation, and loss of talin-FAK interactions all significantly reduce the nuclear localization of YAP and the transcriptional activity of YAP^48^. Clearly, focal adhesions and their core components (talin, vinculin, and FAK) are very important regulators of YAP. However, our experiments here indicate that this regulation is not directly regulated by cellular spread area, actin branching based polymerization, or the force acting on talin molecules. An additional important consideration is the dimensionality of the cellular microenvironment. Here, we have focused on mechanosensing in a 2D environment where cells are free to spread and flatten on the surface. However, studies in 3D have shown that activation of YAP can be hindered by an environment that does not allow for cellular penetration into the surroundings and adhesion engagement with the matrix^50^. In these 3D environments, it is likely that actin branching and Arp2/3 may play a role in this process, but additional investigations in 3D environments would be necessary.

In summary, this work identifies an important distinction between the separate mechanosensitive elements within the cell and their direct or indirect impact on YAP mechanosensing. Going forward, it will be important not to conflate these two interdependent physical regulators of YAP (integrin-based adhesions vs. nuclear envelope connections).

Integrin-based adhesions and their force sensitive molecules are essential for sensing of environmental cues, but it is their impact on network level forces that is most critical for mechanosensing through YAP.

## Acknowledgements

The authors would like to thank Dr. James Bear and Dr. Elizabeth Haynes for the gift of the GFP labeled GMF-β over expression construct^33^.

## Ethical Approval

Not Applicable

## Consent to Participate

Not Applicable

## Consent to Publish

All authors reviewed and approved the manuscript.

## Data Availability Statement

All data generated or analyzed during this study are included in this published article and its supplementary information files. Scripts for data analysis and the processed data spreadsheet are available on GitHub (https://github.com/TristanDriscoll/Sadeghifar-Villalobos).

## Author Contributions

A.S. and C.V. equally contributed to performing experiments, analyzing data, writing and editing the manuscript. J.M. - experiments and data analysis. J.D. – PCR experiments and data analysis. G.M. - software for data analysis, writing, and editing of the manuscript. T.D. – conceptualization, methodology, software, writing and editing of the manuscript. All authors reviewed and approved the manuscript.

## Supplemental Figures

**Supplemental Figure 1:**
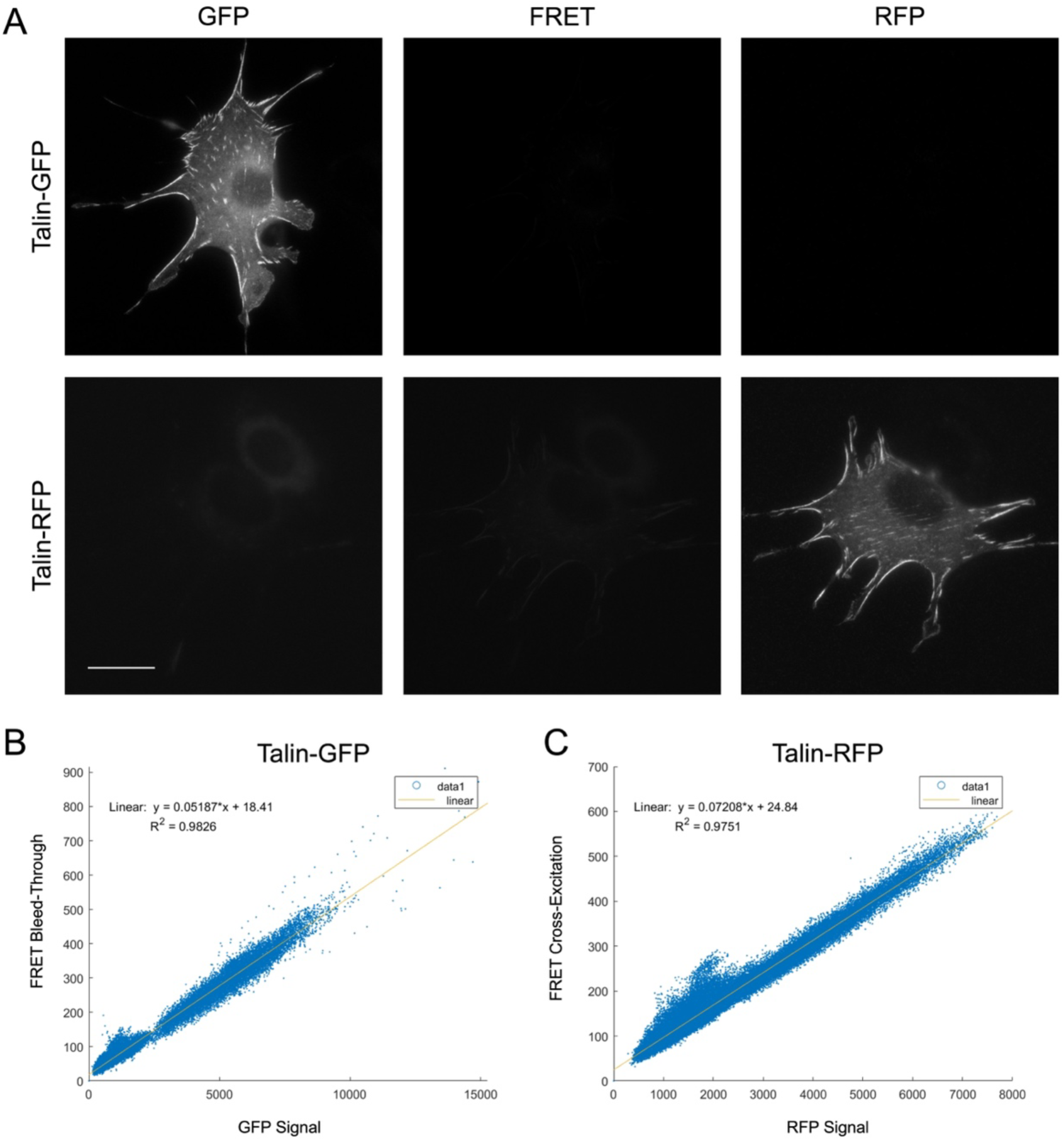
Calculation of the bleed through and cross-excitation coefficients for FRET tension sensor imaging using a GFP tagged Talin (Talin-GFP) and an RFP tagged Talin (Talin-RFP). Example images of each construct expressed in 3T3 cells with each of the 3 channels captured for 3-image FRET calculations (A). Plots of GFP bleed through (B) and RFP cross-excitation (C) with linear fits to extract the bleed through and cross-excitation coefficients for the specific microscope settings used. Scale bar = 20μm.

**Supplemental Figure 2:**
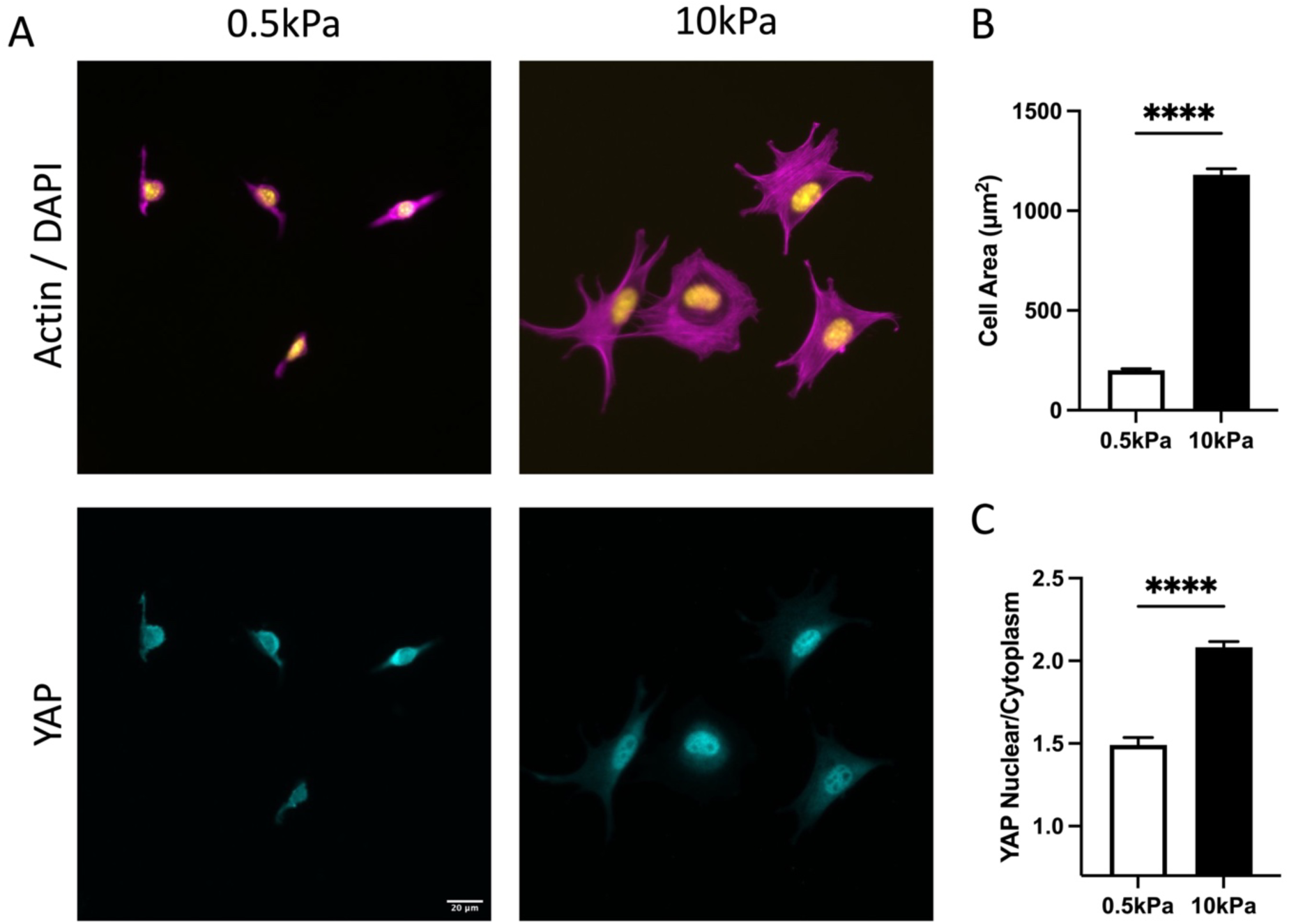
Low stiffness control using a 0.5kPA polyacrylamide gel coated with fibronectin and compared to the 10kPA PDMS gel used for TFM and other deformable substrate experiments. (A) Representative images of Actin/DAPI (magenta/yellow) and YAP (cyan). Quantification of cell area (B) and nuclear to cytoplasmic ratio of YAP (C). Mean +/-SEM, two-sided t-test, **** p<0.0001, n=65-190 cells per group. Scale bar = 20μm.

**Supplemental Figure 3:**
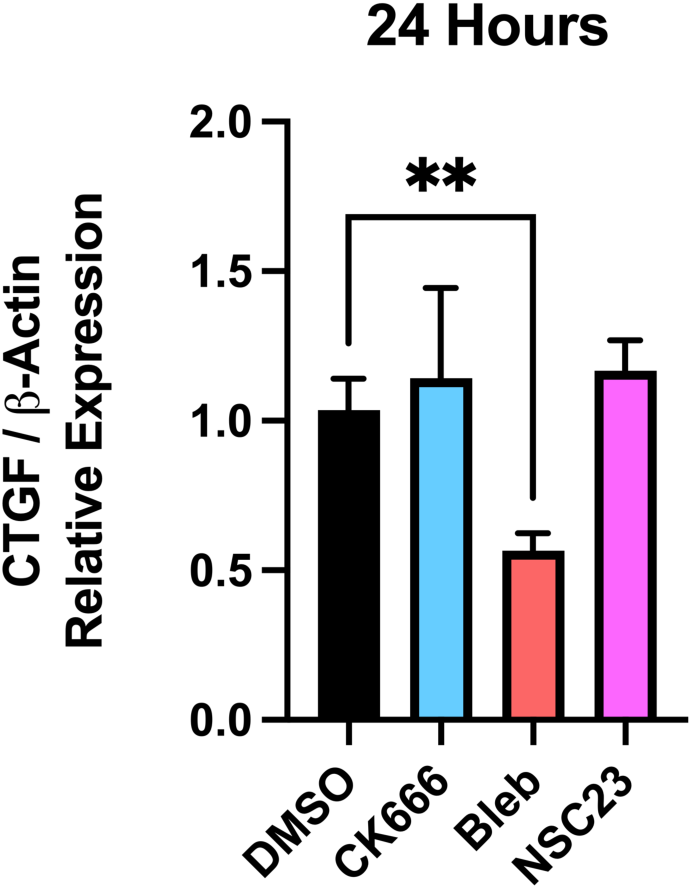
qPCR for expression of the YAP target gene CTGF normalized to expression of β-Actin for cells treated with inhibitors for 24 hours on fibronectin coated glass. DMSO (no inhibitor control), inhibition of Arp2/3 (CK666 50μM), inhibition of myosin (Bleb 10 μM) or inhibition of Rac (NSC23 50μM). Mean +/- SEM, n=3-8 samples per group from 3 independent experiments. One-way ANOVA with Bonferroni’s Post Hoc, ** p<0.01.

**Supplemental Figure 4:**
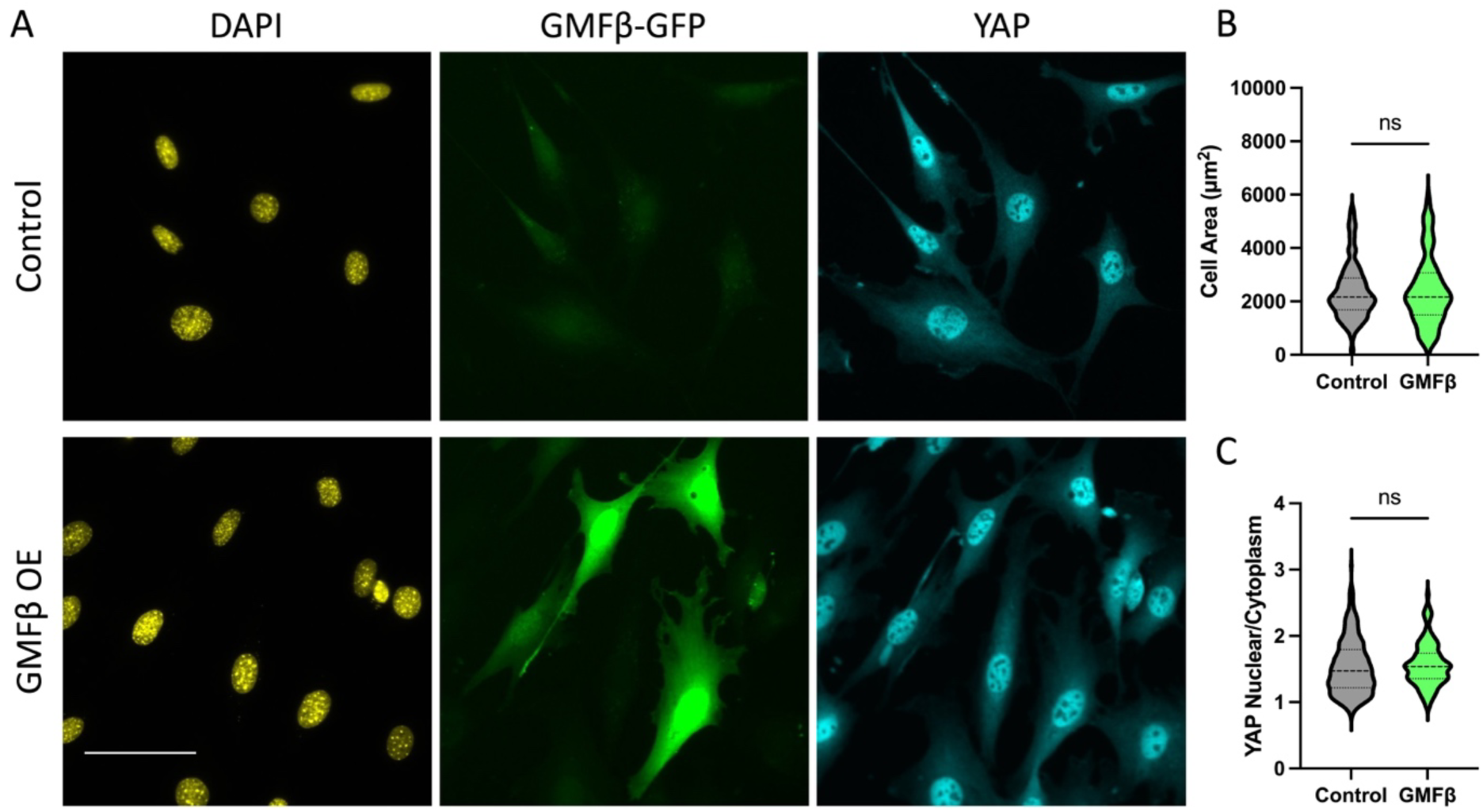
Over expression of the actin debranching protein GMFβ-GFP in 3T3 cells seeded for 24 hours on fibronectin coated glass, with representative images (A) of nucleus (DAPI, yellow), GMFβ-GFP (green), and YAP (cyan). Quantification of cell spread area (B) and YAP nuclear to cytoplasmic ratio (C). Violin plots indicate distribution, mean, and quartiles, n>72 cells per group. Scale bar = 50μm.

**Supplemental Figure 5:**
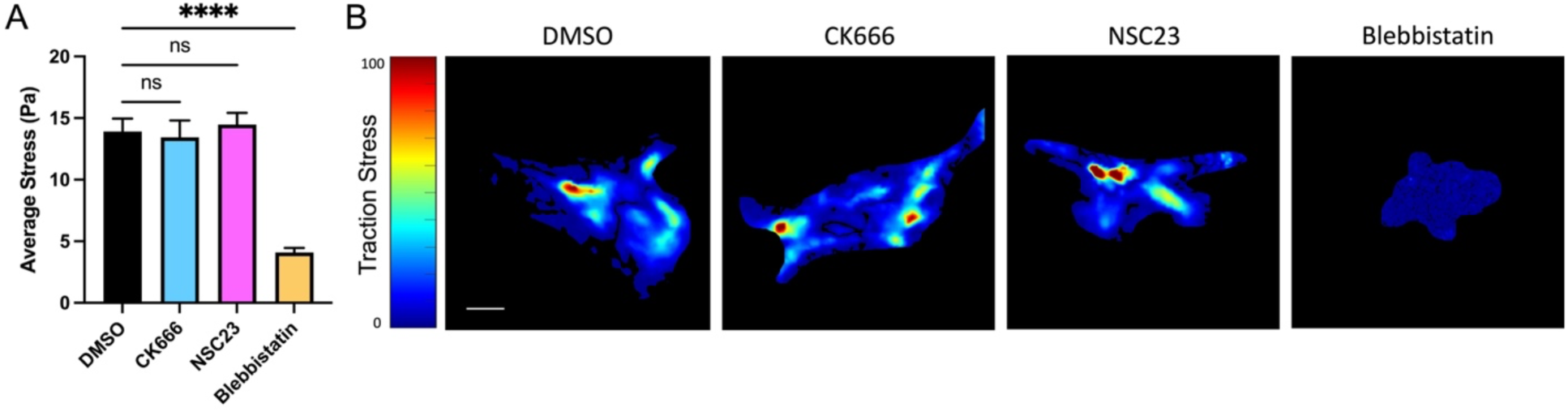
Traction force microscopy on 2kPA PDMS gels coated with fibronectin for 3T3 cells treated with Arp2/3 inhibitor (CK666, 50μM), Rac inhibitor (NSC23, 50μM), or myosin inhibitor (Blebbistatin, 10μM), compared to DMSO control. Quantification of average traction stress per cell (A) and representative heat maps of traction forces (B). Mean +/- SEM, n=12-41 cells per group from 2 independent experiments. Scale bar = 10μm.

## References

1 Kechagia, J. Z., Ivaska, J. & Roca-Cusachs, P. Integrins as biomechanical sensors of the microenvironment. Nat Rev Mol Cell Biol (2019). 10.1038/s41580-019-0134-2

2 Crisp, M. et al. Coupling of the nucleus and cytoplasm: role of the LINC complex. J Cell Biol 172, 41–53 (2006). 10.1083/jcb.200509124

3 Maurer, M. & Lammerding, J. The Driving Force: Nuclear Mechanotransduction in Cellular Function, Fate, and Disease. Annu Rev Biomed Eng 21, 443–468 (2019). 10.1146/annurev-bioeng-060418-052139

4 Kirby, T. J. & Lammerding, J. Emerging views of the nucleus as a cellular mechanosensor. Nature cell biology 20, 373–381 (2018). 10.1038/s41556-018-0038-y

5 Low, B. C. et al. YAP/TAZ as mechanosensors and mechanotransducers in regulating organ size and tumor growth. FEBS Lett 588, 2663–2670 (2014). 10.1016/j.febslet.2014.04.012

6 Zheng, Y. & Pan, D. The Hippo Signaling Pathway in Development and Disease. Dev Cell 50, 264–282 (2019). 10.1016/j.devcel.2019.06.003

7 Gokey, J. J., Patel, S. D. & Kropski, J. A. The Role of Hippo/YAP Signaling in Alveolar Repair and Pulmonary Fibrosis. Front Med (Lausanne) 8, 752316 (2021). 10.3389/fmed.2021.752316

8 Cai, X., Wang, K. C. & Meng, Z. Mechanoregulation of YAP and TAZ in Cellular Homeostasis and Disease Progression. Front Cell Dev Biol 9, 673599 (2021). 10.3389/fcell.2021.673599

9 Atherton, P. et al. Vinculin controls talin engagement with the actomyosin machinery. Nat Commun 6, 10038 (2015). 10.1038/ncomms10038

10 del Rio, A. et al. Stretching single talin rod molecules activates vinculin binding. Science 323, 638–641 (2009). 10.1126/science.1162912

11 Hu, K., Ji, L., Applegate, K. T., Danuser, G. & Waterman-Storer, C. M. Differential transmission of actin motion within focal adhesions. Science 315, 111–115 (2007). 10.1126/science.1135085

12 Thievessen, I. et al. Vinculin-actin interaction couples actin retrograde flow to focal adhesions, but is dispensable for focal adhesion growth. J Cell Biol 202, 163–177 (2013). 10.1083/jcb.201303129

13 Dupont, S. et al. Role of YAP/TAZ in mechanotransduction. Nature 474, 179–183 (2011). 10.1038/nature10137

14 Elosegui-Artola, A. et al. Mechanical regulation of a molecular clutch defines force transmission and transduction in response to matrix rigidity. Nature cell biology 18, 540–548 (2016). 10.1038/ncb3336

15 Chancellor, T. J., Lee, J., Thodeti, C. K. & Lele, T. Actomyosin tension exerted on the nucleus through nesprin-1 connections influences endothelial cell adhesion, migration, and cyclic strain-induced reorientation. Biophysical journal 99, 115–123 (2010). 10.1016/j.bpj.2010.04.011

16 Lele, T. P., Dickinson, R. B. & Gundersen, G. G. Mechanical principles of nuclear shaping and positioning. J Cell Biol 217, 3330–3342 (2018). 10.1083/jcb.201804052

17 Elosegui-Artola, A. et al. Force Triggers YAP Nuclear Entry by Regulating Transport across Nuclear Pores. Cell 171, 1397–1410 e1314 (2017). 10.1016/j.cell.2017.10.008

18 Cosgrove BD, L. C., Driscoll TP, Tsinman TK, Dai EN, Heo S-J, Dyment NA, Burdick JA, Mauck RL. Nuclear envelope wrinkling predicts mesenchymal progenitor cell mechano-response in 2D and 3D microenvironments. Biomaterials (2021). 10.1016/j.biomaterials.2021.120662

19 Driscoll, T. P., Cosgrove, B. D., Heo, S. J., Shurden, Z. E. & Mauck, R. L. Cytoskeletal to Nuclear Strain Transfer Regulates YAP Signaling in Mesenchymal Stem Cells. Biophysical journal 108, 2783–2793 (2015). 10.1016/j.bpj.2015.05.010

20 Goley, E. D. & Welch, M. D. The ARP2/3 complex: an actin nucleator comes of age. Nat Rev Mol Cell Biol 7, 713–726 (2006). 10.1038/nrm2026

21 Krause, M. & Gautreau, A. Steering cell migration: lamellipodium dynamics and the regulation of directional persistence. Nat Rev Mol Cell Biol 15, 577–590 (2014). 10.1038/nrm3861

22 Skau, C. T. & Waterman, C. M. Specification of Architecture and Function of Actin Structures by Actin Nucleation Factors. Annu Rev Biophys 44, 285–310 (2015). 10.1146/annurev-biophys-060414-034308

23 Courtemanche, N. Mechanisms of formin-mediated actin assembly and dynamics. Biophys Rev 10, 1553–1569 (2018). 10.1007/s12551-018-0468-6

24 Pandit, N. G. et al. Force and phosphate release from Arp2/3 complex promote dissociation of actin filament branches. Proc Natl Acad Sci U S A 117, 13519–13528 (2020). 10.1073/pnas.1911183117

25 Papalazarou, V. & Machesky, L. M. The cell pushes back: The Arp2/3 complex is a key orchestrator of cellular responses to environmental forces. Curr Opin Cell Biol 68, 37–44 (2021). 10.1016/j.ceb.2020.08.012

26 Driscoll, T. P. et al. Integrin-based mechanosensing through conformational deformation. Biophysical journal 120, 4349–4359 (2021). 10.1016/j.bpj.2021.09.010

27 Naha, A. & Driscoll, T. P. Fibronectin sensitizes activation of contractility, YAP, and NF-kappaB in nucleus pulposus cells. J Orthop Res 42, 434–442 (2024). 10.1002/jor.25670

28 Gutierrez, E. & Groisman, A. Measurements of elastic moduli of silicone gel substrates with a microfluidic device. PLoS One 6, e25534 (2011). 10.1371/journal.pone.0025534

29 Aratyn-Schaus, Y., Oakes, P. W., Stricker, J., Winter, S. P. & Gardel, M. L. Preparation of complaint matrices for quantifying cellular contraction. J Vis Exp (2010). 10.3791/2173

30 Moro, A. et al. MicroRNA-dependent regulation of biomechanical genes establishes tissue stiffness homeostasis. Nature cell biology 21, 348–358 (2019). 10.1038/s41556-019-0272-y

31 Han, S. J., Oak, Y., Groisman, A. & Danuser, G. Traction microscopy to identify force modulation in subresolution adhesions. Nat Methods 12, 653–656 (2015). 10.1038/nmeth.3430

32 Jang, J. W. et al. RAC-LATS1/2 signaling regulates YAP activity by switching between the YAP-binding partners TEAD4 and RUNX3. Oncogene 36, 999–1011 (2017). 10.1038/onc.2016.266

33 Haynes, E. M. et al. GMFbeta controls branched actin content and lamellipodial retraction in fibroblasts. J Cell Biol 209, 803–812 (2015). 10.1083/jcb.201501094

34 Kumar, A. et al. Talin tension sensor reveals novel features of focal adhesion force transmission and mechanosensitivity. J Cell Biol 213, 371–383 (2016). 10.1083/jcb.201510012

35 Driscoll, T. P., Ahn, S. J., Huang, B., Kumar, A. & Schwartz, M. A. Actin flow-dependent and - independent force transmission through integrins. Proc Natl Acad Sci U S A 117, 32413–32422 (2020). 10.1073/pnas.2010292117

36 Oakes, P. W. et al. Lamellipodium is a myosin-independent mechanosensor. Proc Natl Acad Sci U S A 115, 2646–2651 (2018). 10.1073/pnas.1715869115

37 Justice, R. W., Zilian, O., Woods, D. F., Noll, M. & Bryant, P. J. The Drosophila tumor suppressor gene warts encodes a homolog of human myotonic dystrophy kinase and is required for the control of cell shape and proliferation. Genes Dev 9, 534–546 (1995). 10.1101/gad.9.5.534

38 Xu, T., Wang, W., Zhang, S., Stewart, R. A. & Yu, W. Identifying tumor suppressors in genetic mosaics: the Drosophila lats gene encodes a putative protein kinase. Development 121, 1053–1063 (1995). 10.1242/dev.121.4.1053

39 Zhu, L., Plow, E. F. & Qin, J. Initiation of focal adhesion assembly by talin and kindlin: A dynamic view. Protein Sci 30, 531–542 (2021). 10.1002/pro.4014

40 Klapholz, B. & Brown, N. H. Talin - the master of integrin adhesions. J Cell Sci 130, 2435–2446 (2017). 10.1242/jcs.190991

41 Zhang, X. et al. Talin depletion reveals independence of initial cell spreading from integrin activation and traction. Nature cell biology 10, 1062–1068 (2008). 10.1038/ncb1765

42 Rahikainen, R., Ohman, T., Turkki, P., Varjosalo, M. & Hytonen, V. P. Talin-mediated force transmission and talin rod domain unfolding independently regulate adhesion signaling. J Cell Sci 132 (2019). 10.1242/jcs.226514

43 Nardone, G. et al. YAP regulates cell mechanics by controlling focal adhesion assembly. Nat Commun 8, 15321 (2017). 10.1038/ncomms15321

44 Plouffe, S. W. et al. The Hippo pathway effector proteins YAP and TAZ have both distinct and overlapping functions in the cell. J Biol Chem 293, 11230–11240 (2018). 10.1074/jbc.RA118.002715

45 Das, A., Fischer, R. S., Pan, D. & Waterman, C. M. YAP Nuclear Localization in the Absence of Cell-Cell Contact Is Mediated by a Filamentous Actin-dependent, Myosin II-and Phospho-YAP-independent Pathway during Extracellular Matrix Mechanosensing. J Biol Chem 291, 6096–6110 (2016). 10.1074/jbc.M115.708313

46 Aragona, M. et al. A mechanical checkpoint controls multicellular growth through YAP/TAZ regulation by actin-processing factors. Cell 154, 1047–1059 (2013). 10.1016/j.cell.2013.07.042

47 DeMali, K. A., Barlow, C. A. & Burridge, K. Recruitment of the Arp2/3 complex to vinculin: coupling membrane protrusion to matrix adhesion. J Cell Biol 159, 881–891 (2002). 10.1083/jcb.200206043

48 Holland, E. N. et al. FAK, vinculin, and talin control mechanosensitive YAP nuclear localization. Biomaterials 308, 122542 (2024). 10.1016/j.biomaterials.2024.122542

49 Lachowski, D. et al. FAK controls the mechanical activation of YAP, a transcriptional regulator required for durotaxis. FASEB J 32, 1099–1107 (2018). 10.1096/fj.201700721R

50 Caliari, S. R., Vega, S. L., Kwon, M., Soulas, E. M. & Burdick, J. A. Dimensionality and spreading influence MSC YAP/TAZ signaling in hydrogel environments. Biomaterials 103, 314–323 (2016). 10.1016/j.biomaterials.2016.06.061

